# Evaluating the antiviral activity of Termin-8 and Finio against a surrogate ASFV-like algal virus

**DOI:** 10.1101/2025.05.14.654024

**Authors:** Amanda Palowski, Francisco Domingues, Othmar Lopez, Nicole Holcombe, Gerald Shurson, Declan C. Schroeder

**Affiliations:** Department of Veterinary Population Medicine, College of Veterinary Medicine, University of Minnesota, St. Paul, MN, USA; Anitox Corporation, Lawrenceville, GA, USA; Department of Animal Science, College of Food Agricultural and Natural Resource Sciences, University of Minnesota, St. Paul, MN, USA

**Author notes:** **Correspondence:** Declan C Schroeder.

**Keywords:** African swine fever virus, Emiliania huxleyi virus, NCLDVs, viral inactivation, viability PCR

## Abstract

**Introduction:** Anitox has developed and markets Termin-8 (a formaldehyde-based product) and Finio (non-formaldehyde solution) for the control microbial contamination in feed. To date, both Finio and Termin-8 have not been tested for their potential antiviral activity against megaviruses, such African swine fever virus (ASFV) and its surrogate algal virus, Emiliania huxleyi virus (EhV). Given the limited access and great expense for routine chemical mitigation testing with ASFV, we focused on using EhV as a safe and effective surrogate to evaluate the antiviral activity of these Anitox products. The specific objective of the current study was to evaluate the time course of incubation from hours to days to mimic possible field relevant exposure times for the potential preventative mitigation of megaviruses using Termin-8 and Finio.

**Methods:** Emiliania huxleyi virus was treated with the Anitox recommended concentrations of Termin-8 (0.1% to 0.3% final) and Finio (0.05% to 0.2% final) in biological triplicate experiments. Both viability qPCR (V-qPCR) and standard PCR (S-qPCR) were conducted for EhV copies at 1 hr, 5 hrs, 24 hrs and day 7 post-inoculation.

**Results:** We observed that both Termin-8 and Finio, at their highest treatment concentrations, showed the greatest log reduction of 2 and 4.5 log_10_ units, respectively, at the earliest 1 hr post-inoculation time point. Although Termin-8 efficacy did not improve with time, treatment with Finio showed 100% viable viral inactivation (>5 log_10_ reduction units) at the lowest concentration after 7 days of exposure. In addition, Finio showed negligible viral DNA removal post-treatment as observed via S-qPCR, but use of Termin-8 resulted in a reduction of viral DNA (S-qPCR) equivalent to that observed using the V-qPCR assay.

**Discussion:** Our results demonstrate for the first time that both Termin-8 and Finio can be used as effective chemical mitigants against megaviruses such as EhV and ASFV. Moreover, the mechanism of chemical mitigation involves degradation of the virus particle itself. However, although Termin-8 efficacy appeared to be less than that of Finio, this is likely not the case because Termin-8 cross-links protein viral capsids and fixes the viral particle to make it non-infectious. With the threat of ASFV introduction into the North American swine industry and other non-infected regions around the world, the use of both Termin-8 and Finio as an effective preventive or mitigation strategy to prevent the transmission of ASFV by reducing particle viability in contaminated feed. Additional research is warranted with ASFV, in the presence of various types of feed ingredients and complete feeds and use bioassays to evaluate infectivity because results from V-qPCR revealed that intact viable particles remain post-treatment.

## 1 Introduction

The United States livestock and poultry industries are under constant threat of transmission of foreign animal diseases. Arguably, the most concerning foreign animal disease threat to the U.S. is African swine fever virus (ASFV), which is a double stranded DNA megavirus [1,2]. African swine fever virus is of particular concern because it is stable in a variety of environments (such as feed ingredients) and at various temperatures [1,2,3,4,5]. Because there is no surveillance or monitoring system to determine viral presence or concentration in the global feed supply chain, there is the possibility that ASFV contaminated feed ingredients could be imported into the U.S. from ASFV positive countries. Furthermore, because of the high thermal resilience of ASFV and the detrimental effects of high thermal processing temperatures on the nutritional value (e.g., reduced amino acid digestibility, vitamin potency loss), the use of chemical mitigants to inactivate ASFV in contaminated free ingredients is emerging as a viable biosecurity strategy in global feed supply chains. However, evaluating the efficacy of chemical mitigants against ASFV is difficult because of the very high biosecurity requirements and restricted use in a limited number of approved research laboratories and high cost. Therefore, a safe and effective risk-free *in situ* non-animal (RISNA) assay has been developed that utilizes the algal megavirus (EhV) as a surrogate for ASFV, which can be used in studies to evaluate efficacy of chemical mitigants [3,5]

Both EhV and ASFV are giant double-stranded DNA viruses with similar structural properties [6], functional properties, and inactivation mechanisms, but EhV does not infect humans, animals, or plants making it a safe surrogate virus for ASFV research [1]. The virion of both EhV and ASFV are similar, with both having a nucleoprotein core structure surrounded by an icosahedral capsid and are enclosed by an external lipid envelope [1,3,7]. Additionally, it has recently been shown that both viruses are thermally stable at temperatures up to 80°C for 20 minutes with similar inactivation kinetics when exposed to high temperatures [3, 8]. It is also well documented that porcine viruses, including ASFV, and the algal surrogate EhV, can survive in feed and feed ingredients used in commercial swine diets [1,4,9,10]. Due to all of these similarities, utilizing the algal surrogate EhV in the RISNA assay for trials and experiments to evaluate mitigation strategy effectiveness is very beneficial and informative because it also involves novel analysis assays including viability quantitative PCR, flow cytometry, and confocal microscopy to determine viral presence and structure after treatment [11]. Viability quantitative PCR (V-qPCR) is a useful method to quantify the amount of viable virus in a sample, and viability is defined as an intact virus particle that has the potential to initiate an infection.

Anitox is a global feed additive company that develops a wide range of products to control microbial contamination in feed, such as Termin-8 and Finio. Termin-8 is a formaldehyde-based product with established efficacy against *Salmonella* species [12,13]. Finio was launched in 2017 in the EU and is currently undergoing the registration process in the U.S. as a non-formaldehyde solution that is a patent protected formulation comprised of novel phytochemical and carboxylic acids. To date, both Finio and Termin-8 have not been tested for their potential antiviral activity against megaviruses such ASFV and its surrogate algal virus, EhV. Given the limited access and expense of routine chemical mitigation testing with ASFV, we used the EhV surrogate to evaluate the antiviral effects and efficacy of Termin-8 and Finio in this study. Therefore, the objective of this study was to evaluate the time course of incubation from hours to days to determine the antiviral potential of Termin-8 and Finio as a preventative inactivation treatment of megaviruses, such as EhV and ASFV.

## 2 Materials and Methods

### 2.1 Cell culture and EhV-86 stock

A culture of *Emiliania huxleyi* CCMP374 was grown in Alga-Gro® Seawater Medium (Carolina Biological Supplement Company, NC) at 15°C with 18h/6h light/dark cycle (ca. 2400 lux) until the concentration of 2 × 10^5^ cells/mL was reached. Isolate EhV-86 was added to *E. huxleyi* at a multiplicity of infection of 1 and grown in a 15°C incubator until lysis was observed, which usually occurred after 4 d [7]. The lysate was filtered through a 0.45 µm filter (Nalgene™ Rapid-Flow™ Bottle Top Filters, ThermoFisher Scientific, MA) to remove cell debris. This filtration and infection procedure was repeated several times. The filtered lysate was divided into aliquots and kept in the dark at 4°C until use.

Aliquots of EhV (100 uL containing up to 6 log10 copies/mL) in biological triplicates were treated with manufacturer recommended concentrations of Termin-8 (0.1%, 0.2%, and 0.3% final concentration) and Finio (0.05%, 0.1% and 0.2% final concentration). The EhV triplicates were exposed to varying concentrations of Termin-8 and Finio for 1 hour, 24 hours and 7 days at 20°C.

### 2.2 Standard qPCR and viability qPCR assays

Standard and viability qPCR (S-qPCR and V-qPCR) analyses were conducted for all samples using the methods optimized by Balestreri et al. [3]. For the samples evaluated for virus viability (V) using V-qPCR, 1μL of PMAxx dye (Biotium Inc, CA, 25 µM final concentration) was added and incubated at room temperature in the dark for 10 min on a rocker for optimal mixing. The mixed V samples were exposed to the light using PMA-Lite device (Biotium Inc, CA, US) for 30 minutes to cross-link the PMAxx dye to the DNA. The standard (S) samples analyzed using S-qPCR were not treated with PMAxx dye and were used as a control.

A Nucleo Mag® Virus (Takara Bio USA, Inc.) extraction kit was used to extract DNA from all samples. A quantitative PCR was conducted using Quantinova SYBR Green PCR kit (Qiagen, CA, US) with the following conditions: 2 min at 95°C followed by 40 cycles of 5 s at 95°C and 10 s at 60°C using methods by Balestreri et al. 2024[3]. All PCR assays were conducted using Roto-gene Q Real-Time PCR (Qiagen, CA, US).

### 2.3 Flow cytometry

A 50μL volume of EhV-86 treated with 0.3% (final concentration) Termin-8 and 0.2% (final concentration) Finio and non-treated EhV-86 were sampled and fixed with glutaraldehyde (0.5% final concentration) as described in Balestreri et al. 2024 [3]. All samples were subsequently diluted 1:100 in 1 mL final volume, stained with SYBR gold (1/100,000 final dilution) and analyzed on an Accuri C6 flow cytometer (BD Biosciences, California, USA) using FL1-H threshold at 700 and fast fluidic rate (66μL min^-1^).

### 2.4 Statistical Analysis

All V-qPCR results were analyzed via ANOVA with an independent two-sample t-tests as applicable in Excel (Microsoft Corporation, 2018) for significance differences among treatments. Statistically significant differences between treatments were designated at a significance level of *p* ≤ 0.05.

## 3 Results

### 3.1 Viability of EhV after exposure to Temin-8 and Finio

After 1 hour of treatment with Termin-8, there was a reduction of viral DNA (S-qPCR) equivalent to that observed in the V-qPCR assay, with the greatest reduction of viral DNA occurring at the highest concentration (0.3%, Figure 1A and 1B). While Finio showed negligible viral DNA reduction after an hour of treatment regardless of dose as observed by S-qPCR, when Finio was added at the highest concentration (0.2%), there was complete inactivation (>5 log reduction units) of viable virus (Figure 1C, 1D).

**Figure 1.**
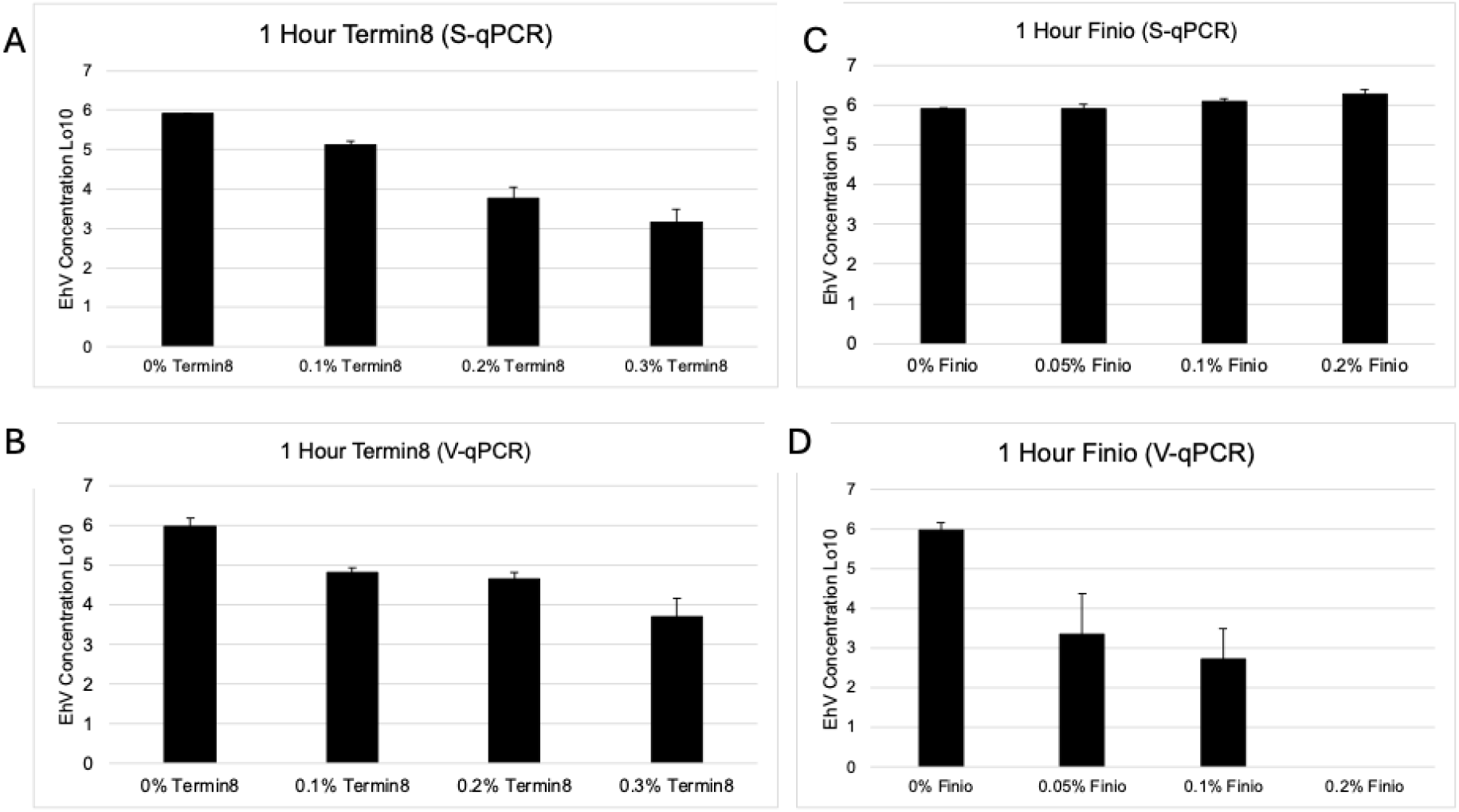
Average log_10_ EhV after 1 hour exposure to 0.1%, 0.2% or 0.3% Termin8 (A & B) or 0.05%, 0.1% or 0.2% Finio (C & D) for S-qPCR (A & C) or V-qPCR (B & D). Bars = standard deviation.

For all concentrations (0.1%, 0.2% and 0.3%) tested, Termin-8 efficacy in viral inactivation improved over time (Figure 2A). At 24 hours, all concentrations (0.1%, 0.2%, and 0.3%) of Termin-8 resulted in significantly lower (*p* = 0.04, 0.05, and 0.05 respectively) concentrations of EhV compared with virus concentrations from 0.1% Termin-8 at 1 hour of exposure. After 7 days incubation at a concentration of 0.1% Termin-8, the amount of viable virus detected via V-qPCR was significantly greater than the 0.3% treatment at 1 hour, and at all concentrations (0.1%, 0.2%, and 0.3%) after 24 hours (*p* = 0.02, 0.01, 0.02, and 0.02 respectively). Furthermore, after 7 days of incubation, the amount of viable viral DNA exposed to 0.2% Termin-8 was significantly greater than the DNA at all concentrations at 1 hour (*p* = 0.05, 0.04, 0.03), all concentrations at 24 hours (*p* = 0.03, 0.03, and 0.03), and with 0.3% Termin-8 at 7 days of incubation (*p* = 0.04).

**Figure 2.**
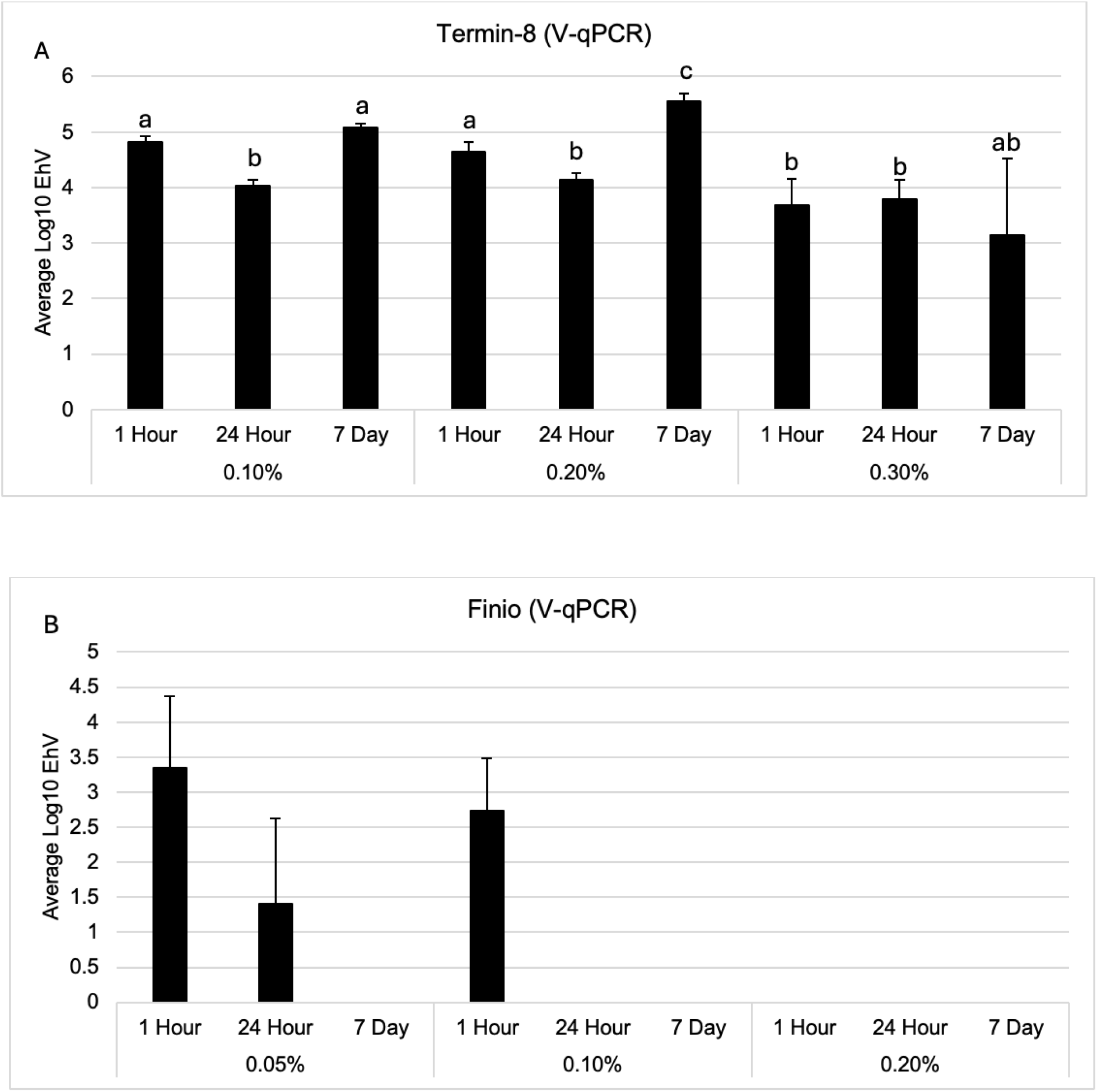
Average viral log_10_ EhV concentrations of Termin-8 (A) and Finio (B) at all time points. Bars = standard deviation. Differing letters denote statistical significance at p ≤ 0.05.

Over time, all tested concentrations (0.05%, 0.1% and 0.2%) of Finio resulted in complete inactivation after 7 days incubation, with a reduction of 5.90 log_10_ units (Figure 2B). Both of the highest dose concentrations (0.3% and 0.2%) of Termin-8 and Finio, respectively, resulted in the greatest log reduction of 2 and 4.5 log_10_ units, respectively, at the earliest (1 hour) timepoint (Figures 1 & 2). At 24 hours, all concentrations (0.1, 0.2, and 0.3%) of Termin-8 had a 1.97, 1.86, and 2.21 log_10_ unit reduction, respectively, but after 7 days of incubation, the lower dosages (0.1% and 0.2%) resulted in less than 1 log_10_ unit reduction (Figure 2A).

All of these results were confirmed by flow cytometry analysis (Figure 3A, 3B, 3C). We found that 0.2% Finio reduced the amount of virus particles detected via flow cytometry (Figure 3B, red rectangle, 8.44 x 10^5^ EhV/mL), while the amount of EhV particles from the 0.3% Termin-8 treatment after 1 hour (Figure 3C, red rectangle, 3.28 x 10^6^ EhV/mL) was similar to the amount of virus in non-treated samples (Figure 3A, red rectangle, 7.45 x 10^7^ EhV/mL).

**Figure 3.**
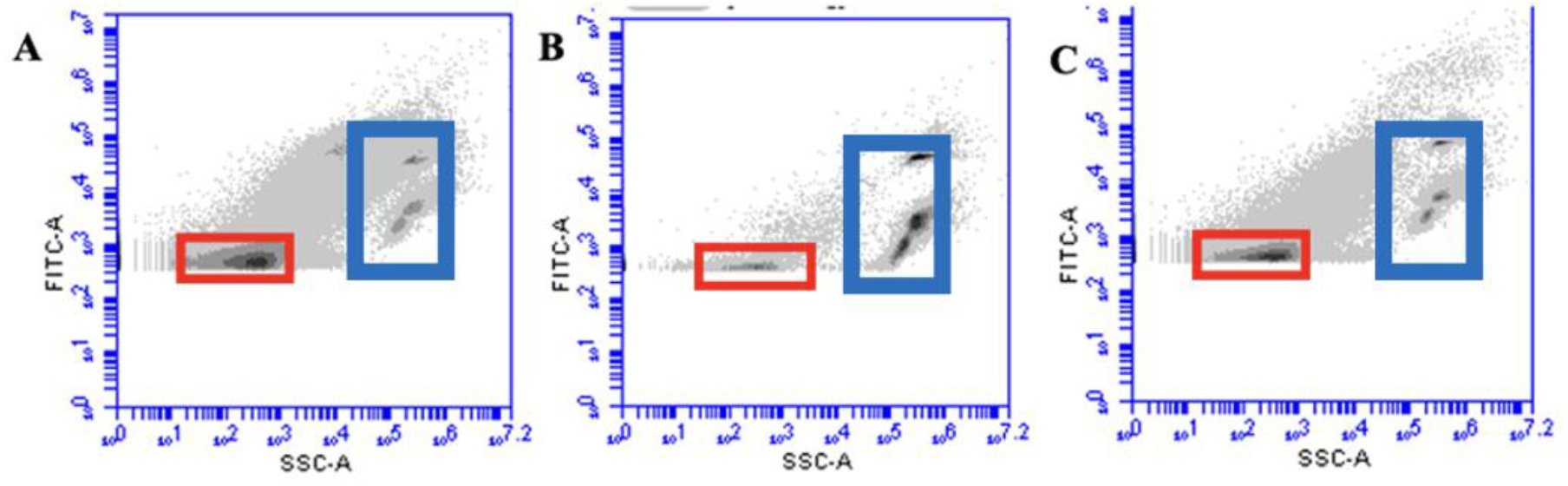
Flow cytometry plots of non-treated (A) and highest concentration of Finio (B) and Termin-8 (C), with red rectangles representing EhV population, while blue rectangles represent control staining beads.

## 4 Discussion & Conclusions

Previous research has shown that megaviruses, such as ASFV and the algal surrogate EhV, are very stable in the environment, particularly at high temperatures [3,5]. Additionally, these viruses have been shown to be relatively stable and survive for up to 21 days in various feed ingredient matrices [1, 2, 4, 5, 8, 9, 10]. Although certain feed ingredients can undergo extremely high temperatures during various processing stages, the amount of time and temperature exposure is not enough to fully inactivate the viral load [3,5]. Therefore, the U.S. feed and swine industry is interested in developing feed supply chain biosecurity protocols that include effective chemical mitigation strategies for swine viruses including ASFV. Results from previous research have shown that viral inactivation by chemical mitigation may leave the host animal better prepared to deal with infection from that particular virus, yet the mechanism remains largely unknown [14, 15]. In this study we used the viability qPCR method to quantify the amount of intact or “viable” viruses left in a sample after exposure to chemical mitigants. Any degraded, damaged, or compromised viral particles were bound by photoreactive dye and unable to amplify during PCR amplification, thereby determining how many viral particles were left intact after chemical mitigation [16,17]. These viability results give us an idea of the potential for the virus to infect a host after chemical mitigation.

Currently, Termin-8 and Finio are commercially used to control microbial contamination in feed. In this study, we tested the antiviral activity of both compounds to show that they can be effective chemical mitigants of ASFV, which provides the swine industry valuable feed biosecurity strategy. Overall, both Termin-8 and Finio have the potential to be effective chemical mitigants against megaviruses, such as ASFV, with the greatest log reduction of 2 (0.3% Termin-8) and 4.5 (0.2% Finio) log_10_ units observed after 1 hour exposure to EhV. Additionally, the mechanism of chemical mitigation by both Temin-8 and Finio occurs by the degradation of the virus particle itself. Although the results from this experiment appeared to indicate that the efficacy of Termin-8 is less than that of Finio, this is likely not the case because Termin-8 appears to cross-link and fix viral particles, making them non-infectious [18]. However, future bioassay-based experiments are needed to confirm this mechanism. Additionally, research in feedstuffs and ASFV to understand the full scope of how these compounds work in real-world scenarios is warranted.

## 6 Author Contributions

DCS designed the study and use of EhV-86 as a surrogate for ASF. AP maintained the *Emiliania huxleyi* and EhV-86 stocks, performed the experiments, and performed the viability qPCRs. DCS & AP wrote the manuscript. DCS designed the V-qPCR experiments. All authors contributed to the article and approved the submitted version.

## 8 Acknowledgments

The authors would like to thank Anitox Corporation for the funding and research opportunity provided by this project.

